# Facilitated event-related power-modulations during transcranial alternating current stimulation (tACS) revealed by concurrent tACS-MEG

**DOI:** 10.1101/249409

**Authors:** Florian H. Kasten, Burkhard Maess, Christoph S. Herrmann

**Author notes:** Corresponding author: Christoph S. Herrmann, Experimental Psychology Lab, Carl von Ossietzky University, Ammerländer Heerstr. 114 ‒ 118, 26129, Oldenburg, Germany.

## Abstract

Non-invasive approaches to modulate oscillatory activity in the brain receive growing popularity in the scientific community. Transcranial alternating current stimulation (tACS) has been shown to modulate neural oscillations in a frequency specific manner. Due to a massive stimulation artifact at the targeted frequency, only little is known about effects of tACS during stimulation. I.e. it remains unclear how the continuous application of tACS affects event-related oscillations during cognitive tasks. Depending on whether tACS merely affects pre‐ or post-stimulus oscillations or both, stimulation can alter patterns of event-related oscillatory dynamics in various directions or may not affect them at all. Thus, knowledge about these directions is crucial to plan, predict and understand outcomes of solely behavioral tACS experiments. Here, a recently proposed procedure to suppress tACS artifacts by projecting MEG data into source space using spatial filtering was utilized to recover event-related power modulations in the alpha band during a mental rotation task. MEG of twenty-five volunteers was continuously recorded. After 10 minutes of baseline measurement, they received either 20 minutes of tACS at individual alpha frequency or sham stimulation. Another 40 minutes of MEG were acquired thereafter. Data were projected into source space and carefully examined for residual artifacts. Results revealed strong facilitation of event-related power modulations in the alpha band during tACS application. Data provide first direct evidence, that tACS does not counteract top-down suppression of intrinsic oscillations, but rather enhances pre-existent power modulations within the range of the individual alpha (=stimulation) frequency.

**Significance:** Transcranial alternating current stimulation (tACS) is increasingly used in cognitive neuroscience to study the causal role of brain oscillations and cognition. However, online effects of tACS so far largely remain a ‘black box’ due to an intense electromagnetic artifact encountered during stimulation. The current study is the first to employ a spatial filtering approach to recover and systematically study event-related oscillatory dynamics during tACS, which can potentially be altered in various directions. TACS facilitated pre-existing patterns of oscillatory dynamics during the employed mental rotation task, but does not counteract or overwrite them. In addition, control analysis and a measure to quantify tACS artifact suppression are provided that can enrich future studies investigating tACS online effects.

## 1 Introduction

Oscillatory activity of neuronal assemblies is an ubiquitous phenomenon in the brain observed within and between different brain structures and across species (Buzsáki, 2006). During the past decades these oscillations have been linked to a variety of brain functions such as memory, perception and cognitive performance (Klimesch, 1999; Basar et al., 2000; Buzsáki, 2006; Klimesch et al., 2007). Traditionally, these relationships were fruitfully investigated using imaging techniques such as Electro‐ or Magnetoencephalography (EEG/MEG). However, in their nature these approaches are correlational and cannot resolve causal relationships between neural oscillations and cognitive processes. The recent (re-)discovery of non-invasive transcranial electrical stimulation (tES) now allows to directly probe these causal relationships (Herrmann et al., 2016b).

The application of oscillatory currents through the scalp by means of transcranial alternating current stimulation (tACS) has been shown to modulate endogenous brain oscillations in a frequency specific manner (Fröhlich and McCormick, 2010; Ozen et al., 2010; Zaehle et al., 2010; Helfrich et al., 2014). While effects of tACS during stimulation have been investigated in animals (Fröhlich and McCormick, 2010; Ozen et al., 2010; Kar et al., 2017) and computational models (Fröhlich and McCormick, 2010; Reato et al., 2010; Ali et al., 2013), studies on tACS effects in humans have so far mostly been restricted to behavioral measures (Marshall et al., 2006; Kar and Krekelberg, 2014; Lustenberger et al., 2015), BOLD-signal effects (Alekseichuk et al., 2016; Cabral-Calderin et al., 2016; Vosskuhl et al., 2016; Violante et al., 2017) and after-effects in the M/EEG (Zaehle et al., 2010; Wach et al., 2013; Neuling et al., 2015; Veniero et al., 2015; Vossen et al., 2015; Kasten et al., 2016). For the later, a frequency specific increase in oscillatory power after stimulation is consistently reported (Zaehle et al., 2010; Neuling et al., 2013; Vossen et al., 2015; Kasten et al., 2016).

Besides outlasting effects on the power of spontaneous oscillations, tACS has more recently been demonstrated to alter event-related oscillatory dynamics in the context of a cognitive task (Kasten and Herrmann, 2017). In that study, event-related desynchronization (ERD) was enhanced after tACS application, accompanied by facilitated performance in a classic mental rotation (MR) task (Shepard and Metzler, 1971; Kasten and Herrmann, 2017). The amount of ERD in the alpha band had previously been linked to MR performance (Michel et al., 1994; Klimesch et al., 2003). Although an increase in task performance was already observed during tACS, the precise oscillatory dynamics during tACS remain unclear (Kasten and Herrmann, 2017). An understanding of the effect of tACS on event-related oscillations is however crucial, given that many tACS-studies solely measure behavior. Depending on whether the stimulation merely affects pre‐ or post-stimulus oscillations or both, tACS may increase, decrease or not effect patterns of ERD/ERS with different behavioral outcomes to be expected. The current study aims to provide a first step towards understanding the effects of tACS on event-related power-modulations during stimulation. To this end, the experiment of Kasten and Herrmann (2017) was repeated in a MEG setup. The application of linearly constrained minimum variance beamforming (LCMV, Van Veen et al., 1997) on MEG recordings has been shown to substantially suppress electromagnetic artifacts encountered during tES (Soekadar et al., 2013; Neuling et al., 2015). Although this approach will never completely remove artifacts from the signal (Noury et al., 2016; Mäkelä et al., 2017; Noury and Siegel, 2017), artifact suppression might still be sufficient to recover changes in event-related dynamics during tACS (Neuling et al., 2017).

Here, this approach was utilized to recover the event-related power-modulations in the alpha band encountered during MR. Based on previous behavioral results, an increase in alpha power-modulation during tACS was hypothesized (Kasten and Herrmann, 2017). Measures to capture tACS effects were carefully chosen to be robust against the possible influence of residual artifacts in the data and control analyses were conducted to rule out that the observed effects can be attributed to a residual artifact.

## 2 Methods

### 2.1 Participants

Twenty-five healthy volunteers were randomly assigned to one out of two experimental conditions. They received either 20 minutes of tACS or sham stimulation throughout the course of the experiment. All were right-handed according to the Edinburgh-handedness scale (Oldfield, 1971) and had normal or corrected to normal vision. Participants gave written informed consent prior to the experiment and reported no history of neurological or psychiatric conditions. The experiment was approved by the *“Commission for Research Impact Assessment and Ethics”* at the University of Oldenburg and conducted in accordance with the declaration of Helsinki. Three subjects exhibited low tolerance to skin or phosphene sensations while determining the individual stimulation intensity (see section 2.3). Due to the resulting low stimulation currents (below 0.4 mA) these subjects were excluded from the analysis. Furthermore, two participants were excluded as they did not exhibit alpha modulation in response to the cognitive task during the baseline block. Data of twenty subjects (10 in stimulation group 10 in sham, age: 26 ± 3 years, 8 females) remained for analysis. Although experiment was initially counterbalanced for participants’ sex, the exclusion of subjects resulted in an imbalance in the sham group (7 males vs. 3 females, 5 males vs. 5 females in the stimulation group).

### 2.2 Magnetoencephalogram

Neuromagnetic activity was recorded at a rate of 1000 Hz using a 306 channel whole-head MEG system (Elekta Neuromag Vectorview, Elekta Oy, Helsinki, Finland) with 102 magnetometers and 204 orthogonal, planar gradiometers, sampling from 102 distinct sensor locations. An online band-pass filter between 0.1 Hz and 330 Hz was applied. The experiment was conducted in a dimly lit, magnetically shielded room (Vacuumschmelze, Hanau, Germany) with participants seated below the MEG helmet in upright position. Three anatomical landmarks (nasion, left and right pre-auricular points), the location of five head position indicator (HPI) coils, as well as > 200 head shape samples were digitized prior to the experiment for continuous head-position tracking and later co-registration with anatomical MRIs using a Polhemus Fastrack (Polhemus, Colchester, VT, USA).

After finishing the preparations, individual alpha frequency (IAF) was determined from a three-minute eyes-open resting state MEG recording. Data were segmented into 1 s epochs. Fast Fourier Transforms (FFTs) were computed for each of the segments using the Fieldtrip toolbox (Oostenveld et al., 2011). The power peak of the averaged spectra in the 8-12 Hz band was visually identified based on a set of posterior sensors showing most pronounced alpha peaks and used as stimulation frequency in the subsequent procedures (see **Figure 1A** for an overview of the time course of the experiment).

**Figure 1:**
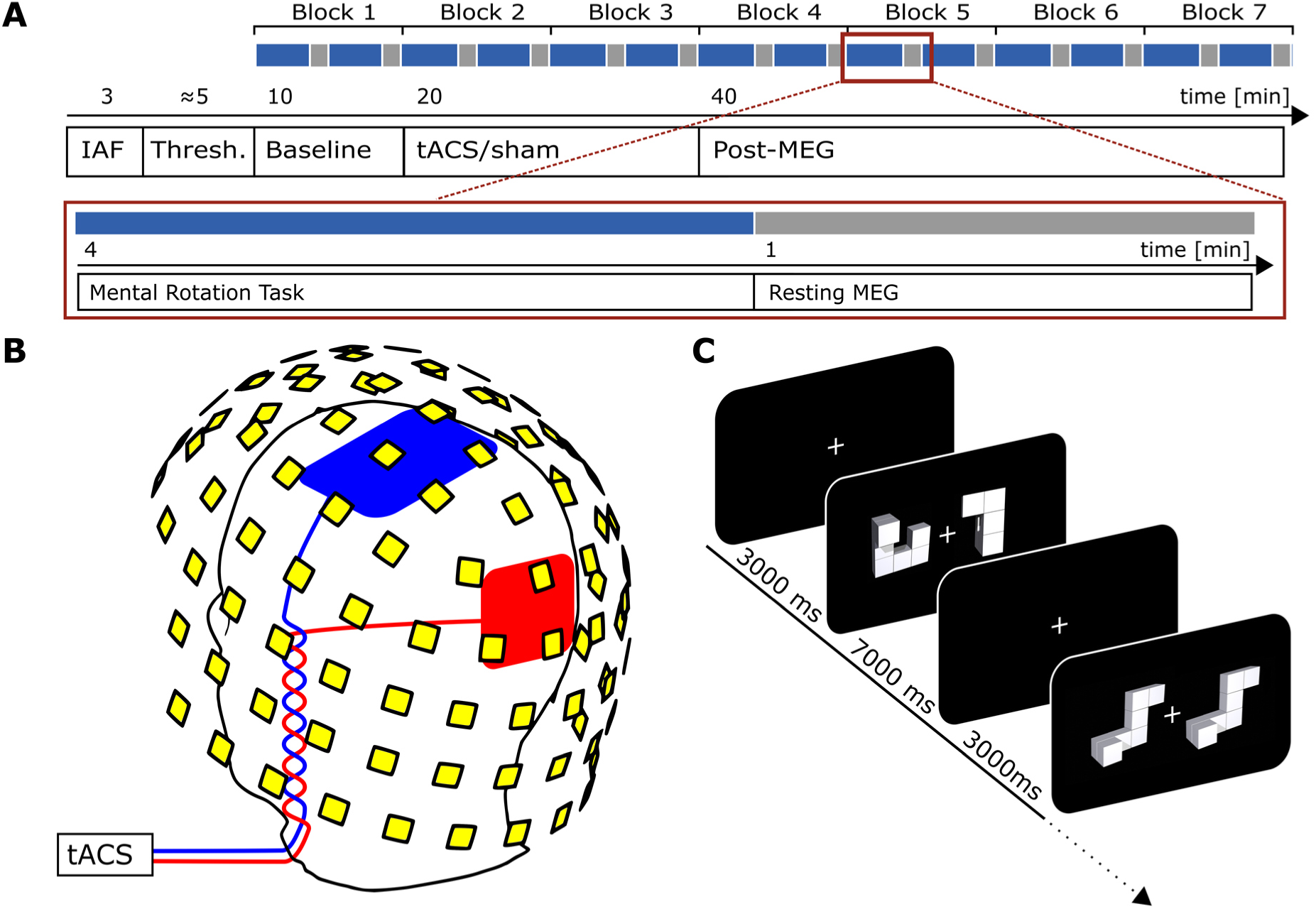
Experimental Procedures. **(A)** Time course of the experiment. Blue color indicates periods during which the MR task was performed, grey indicates intermitting resting periods. **(B)** Positions of stimulation electrodes (red/blue) and layout of MEG sensory (yellow). Stimulation electrodes were placed centered above Cz (7 x 5 cm) and Oz (4 x 4 cm) of the international 10-10 system. MEG was recorded from 102 locations. Each location contains a sensor triplet of one magnetometer and two orthogonal planar gradiometers, resulting in a total of 306 channels. **(C)** Mental rotation task. Each trial started with the presentation of a white fixation cross at the center of the screen. After 3000 ms a mental rotation stimulus was presented and remained on screen for another 7000 ms. During this time participants were required to judge whether the two objects presented were either different (example depicted in 2^nd^ display) or identical (only rotated, 4^th^ display). **A** and **C** are adapted from Kasten and Herrmann (2017).

### 2.3 Electrical stimulation

Participants received either 20 minutes of tACS (including 10 s fade-in and fade-out) or sham stimulation (30 s stimulation in the beginning of the stimulation period, including 10 s fade-in and out) at their individual alpha frequency (IAF). The sinusoidal stimulation signal was digitally generated at a sampling rate of 10 kHz in Matlab 2012a (16-bit, The MathWorks Inc., Natick, MA, USA) and transferred to a digital-analog converter (Ni USB, 6221, National Instruments, Austin, TX, USA). From there the signal was streamed to the remote input of a battery-driven constant current stimulator (DC Stimulator Plus, Neuroconn, Illmenau, Germany), which was placed inside an electrically shielded cabinet outside the MSR. The signal was then gated into the MSR via a tube in the wall using the MRI extension-kit of the stimulator (Neuroconn, Illmenau, Germany). Stimulation was administered by two surface conductive rubber electrodes attached to participants scalp over electrode positions Cz (5 x 7 cm) and Oz (4 x 4 cm) of the international 10-20 system (**Figure 1B**), using an adhesive, electrically conductive paste (ten20 conductive paste, Weaver and Co., USA). Impedance was kept below 20 kΩ (including two 5 kΩ resistors in the cables of the MRI extension-kit of the stimulator). Accordingly, impedance directly under the electrode was limited to 10 kΩ.

To minimize confounding influences from either phosphene or skin sensations, tACS was applied below participants’ individual sensation threshold, using an established thresholding procedure (Neuling et al., 2013, 2015; Kasten et al., 2016; Kasten and Herrmann, 2017). To this end, participants were stimulated with an initial intensity of 500 μA at their IAF. Depending on whether participants noticed the initial stimulation, intensity was either increased or decreased in steps of 100 μA until they noticed/not noticed the stimulation. The highest intensity at which participants did not notice the stimulation was later used as tACS intensity in the main experiment. The thresholding was performed for both groups in order to keep experimental procedures similar. The obtained intensities for the sham group were applied during the 30 s stimulation train in the beginning of the stimulation block (see above). Three participants exhibited sensation threshold below 400 μA and were excluded from analysis. On average, participants were stimulated with 715 μA ± 301 μA (peak-to-peak; stimulation group: 680 μA ± 175 μA) at a frequency of 10.5 Hz ± 0.9 Hz. TACS or sham stimulation was applied for 20 minutes during the second and third block of the behavioral experiment.

### 2.4 Mental rotation task

Visual stimuli were presented using Psychtoolbox 3 (Kleiner et al., 2007) implemented in the same Matlab script that generated the stimulation signal. Visual stimuli were rear-projected onto a screen inside the MSR at a distance of approx. 100 cm from the participant.

Subjects performed the same MR paradigm that was employed in a recent tACS-EEG study (Kasten and Herrmann, 2017). Stimuli were taken from an open-source stimulus set (Ganis and Kievit, 2015), comprised of 384 MR stimuli similar to the objects used in the seminal paper of Shepard and Metzler (1971). The duration of the experiment was reduced from 8 to 7 blocks of 10 minutes each. Participants were familiarized with the task on a notebook during electrode preparation (16 practice trials with immediate feedback). All other parameters were kept similar. Each block consisted of 48 trials, starting with the presentation of a white fixation cross at the center of the screen. After 3000 ms a MR stimulus was presented for another 7000 ms. During this time participants were asked to judge whether the two objects on screen were either identical (can be brought in alignment by rotating) or different (cannot be brought in alignment by rotating) by pressing a button with their left or right index finger (**Figure 1C**). To keep visual stimulation at a constant level, the MR stimuli remained on screen for the whole 7000 ms, regardless of participants’ reaction times. Every 24 trials, the task was interrupted by a one minute resting period during which a rotation of the fixation cross had to be detected. This ensured participants remained focused and tried to avoid head movements. The first block served as baseline measurement before stimulation. During the second and third block, tACS or sham stimulation was applied. The remaining four blocks served as post stimulation measurements to capture aftereffects of the stimulation (**Figure 1A**). The experiment had a total duration of 70 minutes.

### 2.5 Debriefing

After finishing the experiment, participants filled out a translated version of a questionnaire assessing commonly reported side effects of transcranial electrical stimulation (Brunoni et al., 2011). Subsequently, they were asked to indicate whether they believe they received tACS or sham stimulation. Finally, all subjects were informed about the aims of the experiment and their actual experimental condition.

### 2.6 Data analysis

Data analysis was performed using Matlab 2016a (The MathWorks Inc., Natick, MA, USA).

#### 2.6.1 Behavioral data

Analysis of performance and reaction time (RT) data followed the approach of Kasten and Herrmann (2017). Performance in each block (48 Trials) in percent was calculated and normalized by pre-stimulation baseline to account for inter-individual differences. The resulting values reflect performance change in each block relative to baseline. RTs were averaged separately for each rotation angle and normalized by their respective baseline RT. The normalized RTs were then averaged over angles for each block. This procedure accounts for the known increase in RT with larger rotation angles (Shepard and Metzler, 1971).

#### 2.6.2 MEG processing and artifact suppression

MEG data were offline resampled to 250 Hz and filtered between 1 and 40 Hz using a 4^th^ order, two-pass Butterworth filter to approximate zero phase. Data were projected into source space by application of a linearly constrained minimum variance (LCMV) beamformer (Van Veen et al., 1997), a procedure that has been demonstrated to suppress artifacts originating from transcranial electrical stimulation (Soekadar et al., 2013; Neuling et al., 2015). Filter coefficients were individually estimated for each block using the noise covariance matrix, an equally spaced (1.5 cm) 889 point grid warped into Montreal Neurological Institute (MNI) space, and single-shell headmodels (Nolte, 2003), created from individual T1-weighted MRIs. MRIs were co-registered to the median head position in each block, estimated from continuous HPI signals using the Elekta Neuromag MaxFilter™ software (Elekta Oy, Helsinki, Finland). The signal-space-separation method (Taulu et al., 2005), offered by the software was not applied, as it apparently corrupted tACS artefact suppression after beamforming. Covariance matrices were estimated by segmenting each MEG recording into 2 s epochs. The regularization parameter *λ* for the LCMV beamformer was set to zero to ensure optimal artifact suppression as suggested by Neuling et al. (2017).

Sensor space MEG data were segmented ‐5 s to 7 s around the onset of the MR stimuli. Epochs were then projected into sensor space using the previously obtained beamformer filters, resulting in 889 virtual channels, distributed over the brain. A time-frequency analysis was computed for all trials using Morlet-wavelets with a fixed width of 7 cycles. The resulting time-frequency spectra were subsequently averaged for each block.

As mentioned above, all analysis procedures in this study were rigorously checked with respect to their robustness against influences from residual artifacts in the data (Noury et al., 2016; Neuling et al., 2017). This involved a careful choice of the measure used to capture event-related changes in oscillatory power. Traditionally, such changes have been evaluated using the concept of event-related (de-)synchronization (ERD/ERS), which has been defined by Pfurtscheller and Lopes Da Silva (1999) as:

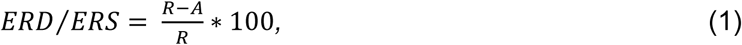

Where *R* is the oscillatory power within the frequency band of interest during a reference period prior to stimulus onset and *A* is the power during a testing period after stimulus onset, respectively. However, assuming that residual tACS artifacts (*R*_*Res*_ and *A*_*Res*_) are equally contributing to *R* and *A*, this would change the equation in the following way:

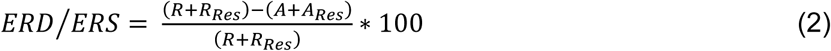

Given that the residuals in *R* and *A* are uncorrelated with the task and have approximately equal strength (*R*_*Res*_ ≈ *A*_*Res*_), their influence cancels out in the numerator, but biases the de-nominator of the equation, resulting in systematic underestimations of the observed power modulations:

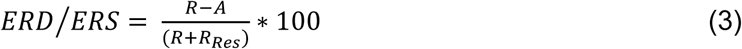

For this reason the pure difference between reference and testing period (for the sake of clarity in the following referred to as event-related power difference; ERΔ_Pow_) was used to more accurately capture event-related power modulations in the current study:

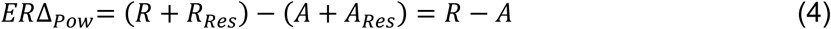

Power in the individual alpha band (IAF ± 2 Hz) was extracted with reference and test period ranging from ‐2.5 s to ‐0.5 before and 0 s to 2 s after stimulus onset, respectively. Performance of the artifact suppression was evaluated by estimating the size of the residual artifact relative to the brain oscillation of interest (see next section). As it will be described in more detail in the results section, the beamformer successfully suppressed the tACS artifact from approx. 2,500,000 times the size of human alpha oscillations down to a factor of < 3. However, some ‘hot spots’ showing larger residual artifacts (1:10) are apparent in the proximity of stimulator cables and the central stimulation electrode. In order to avoid the inclusion of virtual channels in the analysis that contain strong residual artifacts, but no physiologically meaningful effects, brain areas showing strongest alpha power modulation in response to the onset of the MR-stimuli were localized based on the first (artifact-free) block prior to stimulation. To this end a dependent-sample random permutation cluster t-test (two-tailed) with 5000 randomizations and Monte Carlo estimates to calculate p-values was run to compare power in the IAF-band between reference and test period during the baseline block. The test was performed on the whole sample (stimulation and sham group pooled). Clusters were thresholded at an α-level of .01. The resulting significant negative cluster was used as ROI to extract the time course of ERΔ_Pow_ from each block. To account for inter-individual differences, ERΔ_Pow_ in each block was normalized by ERΔ_Pow_ in the baseline block before stimulation. In order to test whether the effects of tACS were specific to the alpha band the same analysis was performed on power modulations in the lower (IAF + 3 Hz to IAF + 11 Hz) and upper (IAF + 12 Hz to IAF + 20 Hz) beta band within the ROI.

##### 2.6.3 Evaluation of artifact suppression and control analyses

As discussed earlier, the application of LCMV beamforming results in a strong, however imperfect suppression of the tACS artifact (Noury et al., 2016; Mäkelä et al., 2017; Noury and Siegel, 2017). It is therefore crucial to characterize the achieved artifact suppression and to rule out that effects observed during stimulation result from residual artifacts in the data rather than an effect of tACS on the brain.

In order to evaluate the artifact suppression achieved by the spatial filtering procedure, participants’ alpha power (IAF ± 2 Hz) was extracted from the pre-stimulus interval of the baseline and the two stimulation blocks. The power in the baseline block provides an estimate of participants’ natural, artifact-free alpha power that can be compared to the power encountered during stimulation blocks before (on the sensor-level) and after beamforming (on the source-level). It is therefore possible to roughly estimate the size of the stimulation artifact relative to the brain signal of interest. This artifact-to-brain-signal-ratio was calculated for each magneto‐ and gradiometer channel as well as for each virtual channel after LCMV. While this measure is not able to disentangle brain signal/tACS effects from a residual artifact after LCMV, it can provide an upper boundary for the size of the residual artifact as well as its spatial distribution.

A major assumption of the presented analysis framework for event-related power modulations during tACS is that the (residual) artifact has similar strength during the pre‐ and poststimulus intervals, such that its influence cancels out when contrasting (subtracting) the two intervals (equation 4). Previous studies have demonstrated that physiological processes such as heartbeat and respiration can result in impedance changes of body tissue and small body movements, which change the size of the tACS artifact (Noury et al., 2016; Noury and Siegel, 2017). To rule out that a similar modulation of artifact strength occurring in an event-related manner accounts for potential effects observed on the source level, a control analysis was carried out. To this end, sensor-level MEG time series during the two stimulation blocks were band-pass filtered around the stimulation frequency (IAF ± 1 Hz) and the signal envelope was extracted using a Hilbert transform. The envelope time-series was subsequently segmented analogously to the ERΔ_Pow_ analysis and demeaned. The difference in envelope amplitude during pre‐ (-2.5s to ‐0.5s) and post-stimulus interval (0 – 2 s) were compared by means of a random permutation cluster t-test with Monte Carlo estimates. To rule out that these differences drive the effects observed on the source-level, the envelope differences were correlated with the ERΔ_Pow_ values obtained earlier. For comparison, the same analysis was performed for the stimulation and sham group. For the sham group, envelope differences should reflect the event-related suppression of alpha power, commonly observed during mental rotation, and therefore highly correlate with the source level ERΔ_Pow_. Pre-/poststimulus envelope differences in the stimulation group, however, should pre-dominantly reflect changes in the tACS artifact. High correlations between sensor-space envelope differences with source level ERΔ_Pow_ would thus indicate that systematic modulations of the tACS artifact drive changes in ERΔ_Pow_, rather than an actual physiological effect of tACS.

##### 2.6.4 Experimental design and statistical analysis

Statistical analysis was realized in a 2 x 6 mixed-effects repeated measures design with the between subject factor *condition* (stimulation vs. sham) and the within subject factor *block* (6 levels). The normalized behavioral (performance, RTs) and physiological (ERΔ_Pow_) data were analyzed using repeated measures ANOVAs (rmANOVA). Greenhouse-Geisser corrected p-values are reported were appropriate. If significant interactions between *condition* and *block* were revealed, analysis was subsequently split into two separate rmANOVAs, one covering effects during stimulation (factors condition: stimulation vs. sham; block: block 2 vs. block 3), the other analyzing outlasting effects (factors condition: stimulation vs. sham; block: block 4 - block 7). Comparisons of single blocks were performed using two-sample t-tests. Generalized *η*^2^ and *Cohen’s d* values are reported as measures of effect size. Pearson correlation coefficients were calculated to relate behavioral and physiological effects, as well as physiological effects and stimulation intensity.

Statistical analysis was performed using R 3.2.3 (The R Core Team, R Foundation for Statistical Computing, Vienna, Austria). Cluster based permutation tests on MEG data were performed in Matlab 2016a using statistical functions implemented in the Fieldtrip toolbox (Oostenveld et al., 2011).

## 3 Results

### 3.1 Behavioral Results

Welch two sample t-test yielded a trend for slightly better raw task performance in the baseline block for the sham group as compared to the stimulation group (*t*_*14.9*_ = ‐2.00, *p* = .06, *d* = .9; *M*_*stim*_ = 87.3%, SD = 3.6%; *M*_*Sham*_ = 91.7%, SD = 5.9%). The rmANOVA on relative performance change revealed a significantly larger facilitation of mental rotation performance relative to baseline in the *stimulation* group as compared to *sham (condition: F*_*1,18*_ = 4.93, *p* = .04, *η*^2^ = 0.14)

Analysis of RTs revealed a trend for the factor *block* (*F*_*5,90*_ = 2.47, *p* = .07, *η*^2^ = 0.03), but no effect of stimulation (*F*_*1,18*_ = 1.02, *p* = .33, *η*^2^ = 0.04). Results of the behavioral analysis are summarized in Figure 2.

**Figure 2:**
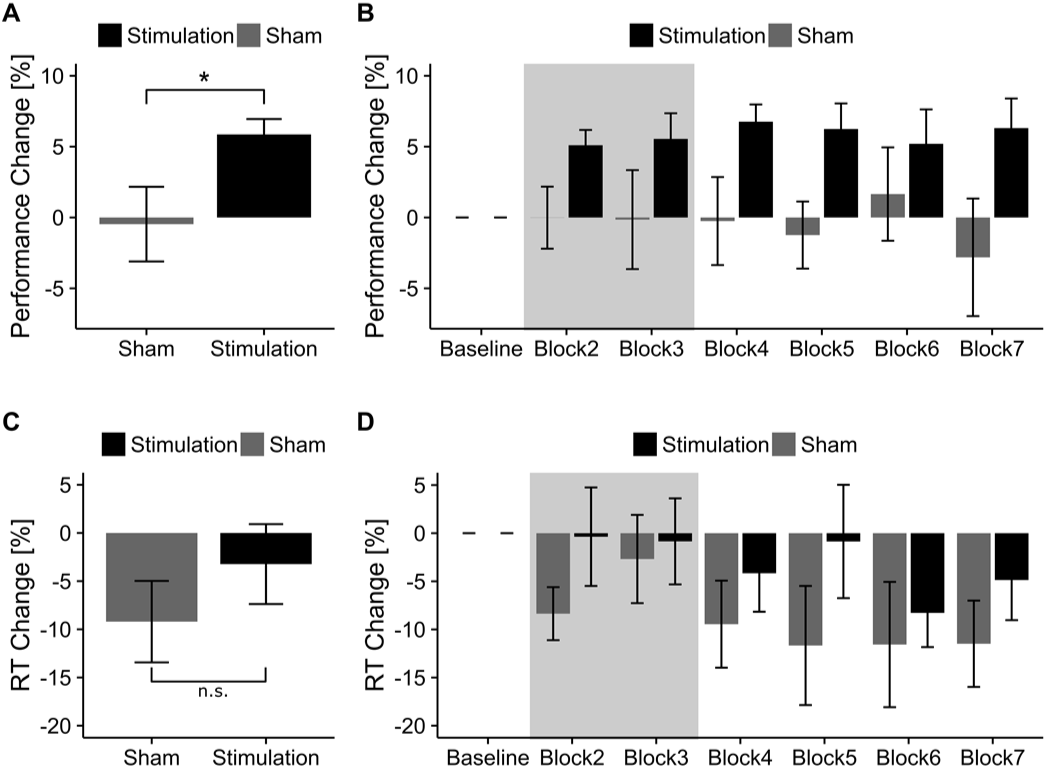
Behavioral results. **(A)** Change in task performance for *stimulation* and *sham* group relative to baseline pooled over all experimental blocks. Error bars depict standard error of the mean (*S.E.M*). Asterisks code for significance (* < .05). **(B)** Change in task performance relative to baseline for *stimulation* and *sham* group depicted over experimental blocks. The grey area indicates blocks that were performed during tACS or sham stimulation. **(C)** Change in RT for *stimulation* and *sham* group relative to baseline pooled over experimental blocks. **(D)** Change in RT for *stimulation* and *sham* group relative to baseline depicted over experimental blocks. Grey area indicates blocks that were performed during tACS or sham stimulation.

### 3.2 Event-related alpha modulation

Comparison of pre‐ and post-stimulus IAF-band power during the baseline block revealed a significant cluster in occipito-parietal areas (*p*_*cluster*_ < .001, **Figure 3A**) for the whole sample. The identified cluster was used as a ROI to extract the time-course of ERΔ_Pow_ from the different blocks and to limit the subsequent analysis to physiologically meaningful brain regions. The subsequent rmANOVA revealed a significant main effect of *block* (*F*_*5,90*_ = 7.22, *p* = .009, *η*^2^ = .15) as well as a significant *condition*block* interaction (*F*_*5,90*_ = 6.81, *p* = .011, *η*^2^ = .15), and a trend for the main effect of *condition* (*F*_*1,18*_ = 3.62, *p* = .07, *η*^2^ = .10). Please refer to **Figure 3B** for an overview of the time course of relative ERΔ_Pow_. To further resolve the significant interaction, separate rmANOVAs were performed on the data acquired during and after stimulation. This analysis exhibited a significant main effect of *condition* (*F*_*1,18*_ = 9.34, *p* = .007, *η*^2^ = .27) during stimulation, but not thereafter (*condition: F*_*1,18*_ = 0.14, *p* = .71, *η*^2^ < .01, **Figure 3C**). Furthermore, a significant effect of *block* (*F*_*3,54*_ = 3.55, *p* = .02, *η*^2^ = .02), as well as a significant *condition*block* interaction (*F*_*3,54*_ = 3.10, *p* = .034, *η*^2^ = .02) were found in the post-stimulation data. None of the other main effects or interactions reached significance. It was not possible to further resolve the significant *condition*block* interaction during the post-stimulation blocks. Separately testing relative ERΔ_Pow_ values of the two experimental groups against each other did not reveal significant differences for any of the blocks (all p > .12, Welch two-sample t-test, one-tailed, uncorrected). Based on pure visual inspection, the effect appears to be driven by a group difference during the first block after stimulation (block 4, see **Figure 3B**), which might be indicative of a weak tACS aftereffect during this block. Refer to **Figure 4** for group-averaged time-frequency representations of participants’ normalized alpha power change and the corresponding source-level topographies within the analyzed ROI.

**Figure 3:**
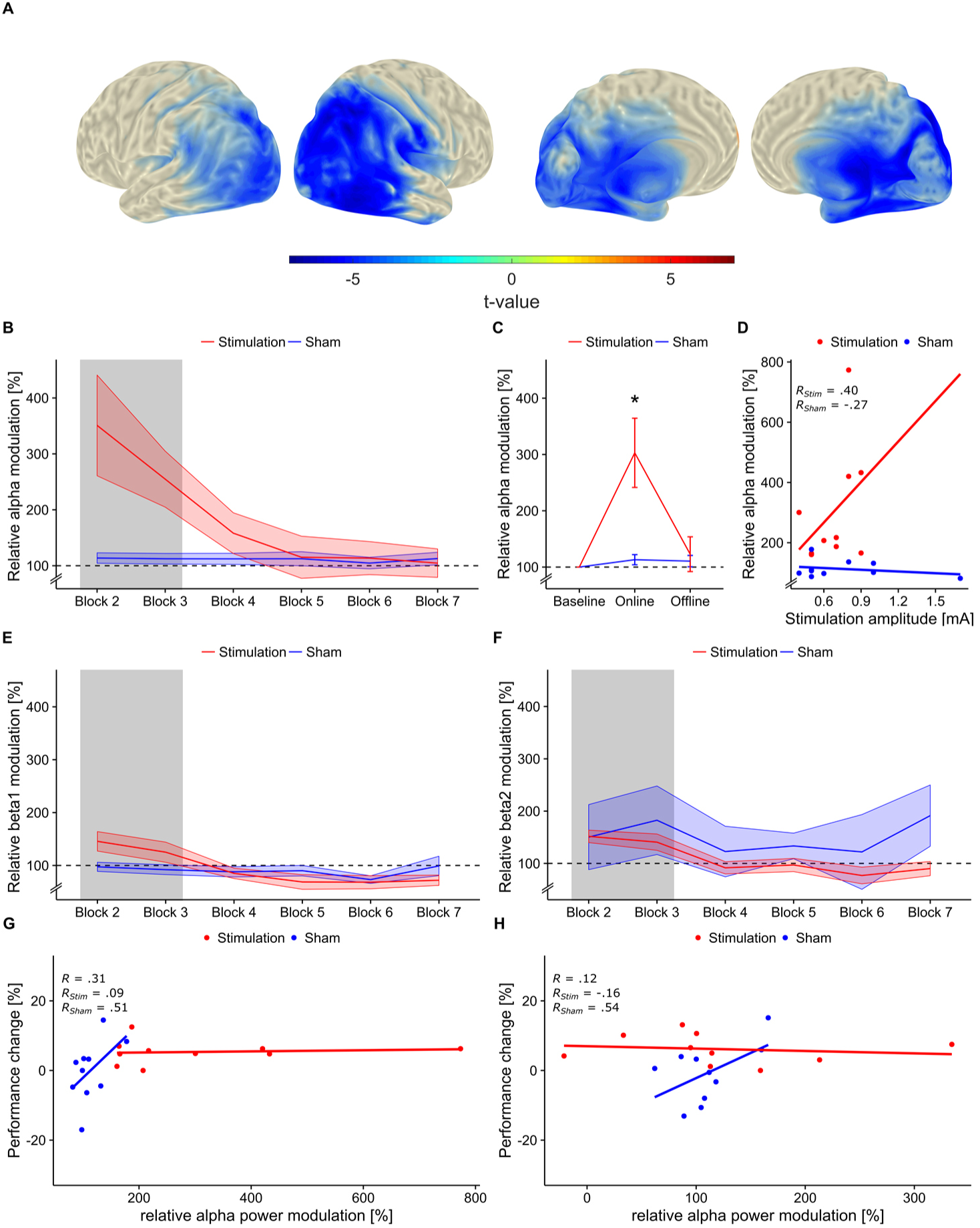
Event-related alpha power modulation. **(A)** Region of interest (ROI). Significant cluster (pre‐ vs. post-stimulus power) in the IAF-band during the first block prior to tACS or sham stimulation, computed pooled on the whole sample (*p*_*cluster*_ < .001). Topographies depict t-values mapped on an MNI standard surface. Statistical maps are thresholded at α < .01. The depicted cluster was used as ROI to extract the time course of alpha power modulation relative to baseline over blocks from the virtual channels. **(B)** Relative alpha power modulation within ROI depicted for each block. The grey area indicates blocks during tACS or sham stimulation. Shaded areas represent standard error of the mean (*S.E.M.*). Dashed line depicts baseline level. **(C)** Relative alpha modulation during tACS or sham (online) and after stimulation (offline). Error bars represent *S.E.M*., asterisks code for significant differences (* < .05). **(D)** Relative alpha modulation during stimulation correlated with stimulation intensity. Each point represents a single subjects’ stimulation amplitude and relative alpha power modulation averaged over the two stimulation blocks (block 2 and 3). Please note that a stimulation intensity was determined for all participants (including sham). However, only for participants in the stimulation group this intensity was continuously applied during block 2 and 3 **(E)** Relative power modulation in the lower beta band (IAF + 3 Hz to IAF + 11 Hz) within the ROI for each block. **(F)** Relative power modulation in the higher beta band (IAF + 12 Hz to IAF + 20 Hz) within the ROI for each block. **(G+H)** Correlation between change in task performance and relative alpha power modulation during **(G)** and after tACS **(H)**. High, albeit non-significant correlations are only evident for the sham, but not the stimulation group.

**Figure 4:**
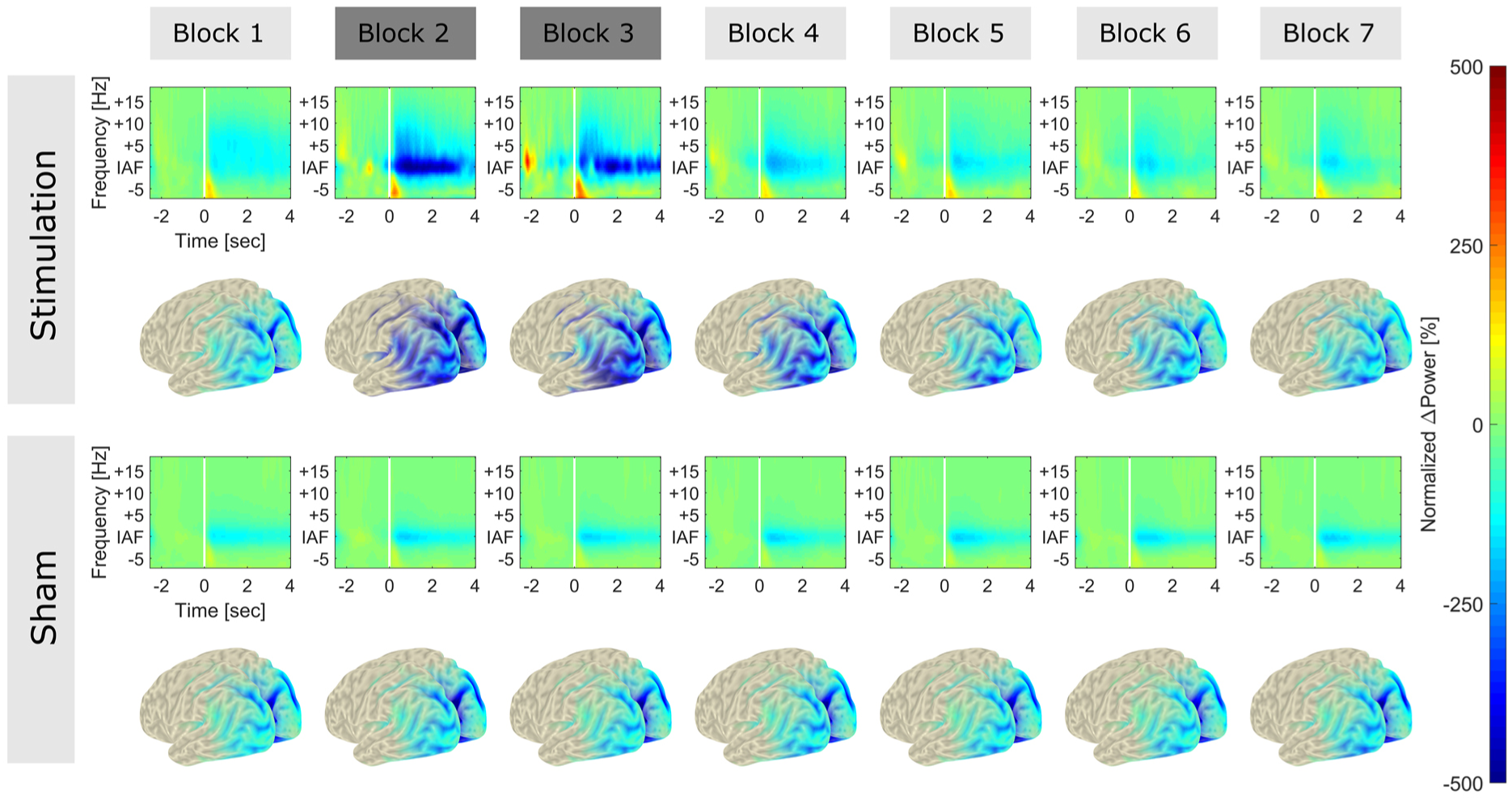
Normalized, baseline-subtracted TFRs and source topographies. TFRs and source topographies for *stimulation* (**Top Rows**) and *sham* group. (**Bottom Rows**). TFRs were aligned at IAF and averaged over subjects in each group. The range from ‐2.5 to ‐0.5 prior to stimulus onset (white bar) served as reference period for baseline subtraction. Spectra were subsequently normalized by the power difference in the alpha band (IAF ± 2Hz) during the baseline block (block 1) prior to stimulation. Normalization was performed such that the data presented resemble data in the statistical analysis. Blocks 2 and 3 (dark grey) represent data acquired during tACS or sham stimulation. All other blocks (light grey) were measured in absence of stimulation. Functional maps were averaged over subjects and projected onto a MNI standard surface. Only activity within the analyzed ROI is depicted. A strong facilitation of event-related power modulation around the IAF can be observed during tACS application (block 2 and 3).

A non-significant positive correlation between the increase in ERΔ_Pow_ during stimulation and stimulation intensity was observed in the *stimulation* group (*r* = .40, *t*_*8*_ = 1.25, *p* = .24). A weak negative non-significant correlation was observed in the *sham* group (*r* = ‐.26, t_8_ = - 0.78, *p* = .45; **Figure 3D**).

To test whether the effects of tACS were specific to the alpha band, the analysis was repeated on event-related power modulations in the lower (IAF + 3 Hz to IAF + 11 Hz) and upper (IAF + 12 Hz to IAF + 20 Hz) beta band within the ROI. The rmANOVA for the lower beta band revealed a significant effect of *block* (*F*_*5,90*_ = 15.10, *p* < .001, *η*^2^ = .17) as well as a significant *condition*block* interaction (*F*_*5,90*_ = 9.37, *p* < .001, *η*^2^ = .11). Two separate rmANOVAs testing the effects during and after stimulation, revealed a trend for the factor *condition* during stimulation (*F*_*1,18*_ = 4.17, *p* = .056, *η*^2^ = .18) as well as a significant effect of *block* (*F*_*1,18*_ = 4.72, *p* = .043, *η*^2^ = .02). After stimulation only a trend for the factor *block* was found (*F*_*3,54*_ = 2.28, *p* = .09, *η*^2^ = .03). No significant effects were found in the analysis of the upper beta band. **Figure 3E,F** summarize results for the lower and upper beta band analysis (all p > .1). There were no significant correlations between relative ERΔ_Pow_ and change in task performance during (*r*_*online*_ = .3, *t*_*18*_ = 1.37, *p* = .18) or after stimulation (*r*_*offline*_ = .11, *t*_*18*_ = 0.49, *p* = .62). Descriptively the correlation was higher for the sham group during both during and after stimulation (*r*_*Sham/online*_ = .51, *t*_*8*_ = 1.67, *p* = .13; *r*_*Sham/offline*_ = .54, *t*_*8*_ = 1.83, *p* = .1) as compared to the stimulation group (*r*_*Stim/online*_ = .09, *t*_*8*_ = 0.27, *p* = .8; *r*_*Stim/offline*_ = ‐.16, *t*_*8*_ = ‐0.45, *p* = .67; **Figure 3G,H**).

### 3.3 Control Analyses

To rule that the strikingly strong facilitation of power-modulations in the alpha band is driven by residual artifacts, some control analyses were performed. In a first step, the performance of the artifact suppression achieved by LCMV was evaluated. To this end, the ratio of pre-stimulus alpha power during the (tACS-free) baseline block and the two tACS blocks was compared in sensor and source space. In the sensor-space, this artifact-to-brain-signal-ratio was on average 2,534,000:1 in block 2 and 2,569,000:1 in block 3 (average over all sensors and subjects). After LCMV beamforming the ratio was reduced to 2.72:1 in block 2 and 3.13:1 in block 3 (average over virtual sensors and subjects). The largest ratio observed in a single virtual channel of one subject after beamforming was 93.42:1. **Figure 5** illustrates the spatial distribution of the alpha to artifact ratio on the source level. The ratio is highest in central areas, covered by stimulation electrodes and cables. Outside these areas the ratio is substantially smaller and falls within a physiologically plausible range for alpha band oscillations (< 3:1/4:1). Overall artifact suppression appears to be slightly worse during block 3 as compared to block 2.

**Figure 5:**
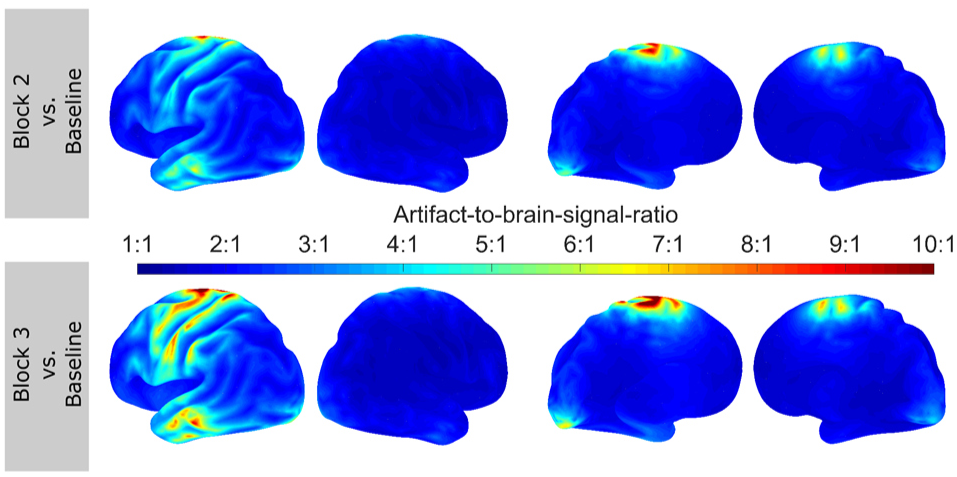
Artifact-to-brain-signal topographies. Topographies depict the average ratio between participants’ pre-stimulus alpha power estimated during the baseline block and residual artifact in the pre-stimulus interval during block 2 (**top row**) and 3 (**bottom row**). Results are depicted only for the stimulation group. The ratio is strongest in central areas covered by stimulation electrodes and cables. Frontal and posterior areas within the ROI seem less affected. Here, the ratio falls in a physiologically plausible range (< 1:3/1:4), such that residual artifact and facilitatory effects of the stimulation or spontaneous increase of alpha power cannot be disentangled. Results have to be interpreted in terms of an upper boundary for the size of the residual artifact, as each virtual channel contains a mixture of brain signal of interest and artifact.

The event-related envelope of the sham group reflects the typical pattern of alpha power decrease after stimulus onset in the MR task in both sensor types. This was supported by the permutation cluster analysis, which revealed significant positive clusters in the magnetometer and the gradiometer data (*p*_*cluster*_ < .001, **Figure 6A,C**; significant sensors are marked by black dots). This was further supported by the high correlation between source-level power modulation and envelope difference of magnetometer (*r* = .96, *t*_*8*_ = 10.17, *p* < .001, **Figure 6B**) and gradiometer channels (*r* = .88, *t*_*8*_ = 5.23, *p* < .001; **Figure 6D**). In the stimulation group time-course and topography of the envelope overall exhibits a reversed pattern with lower amplitudes before stimulus onset and increased amplitude thereafter. In addition, the envelope time-course of gradiometers shows a prominent rhythmic activity in the range of 1 to 2 Hz. Such modulations could potentially reflecting heart-beat related modulations (Noury et al., 2016). However, given that this rhythmic activity was only observed in one sensor type and in a relatively systematic manner, might be more likely to reflect a technical artifact. Importantly, no such rhythmic modulation was evident in the time-frequency representations after LCMV (**Figure 4**). Results of the cluster analysis revealed positive clusters in the gradiometer data in only a few frontal sensors (*p*_*cluster*_ < .05, **Figure 6**, top left) as well as positive and negative clusters for some magnetometer channels (*p*_*cluster*_ < .05). No significant correlation between the observed source-level power modulations and the sensor level envelope differences in magnetometer (*r* = .13, *t*_*8*_ = 0.37, *p* = .72) or gradiometer sensors (*r* = .26, *t*_*8*_ = 0.75, *p* = .47) was evident. Overall, results do not support the idea that the effects observed on the source level can be explained by systematic, task-related changes in artifact strength. Only very few channels exhibit significant task-related power modulations, which, in addition, rather seem so show a reversed pattern of artifact modulation as compared to the source level data.

**Figure 6:**
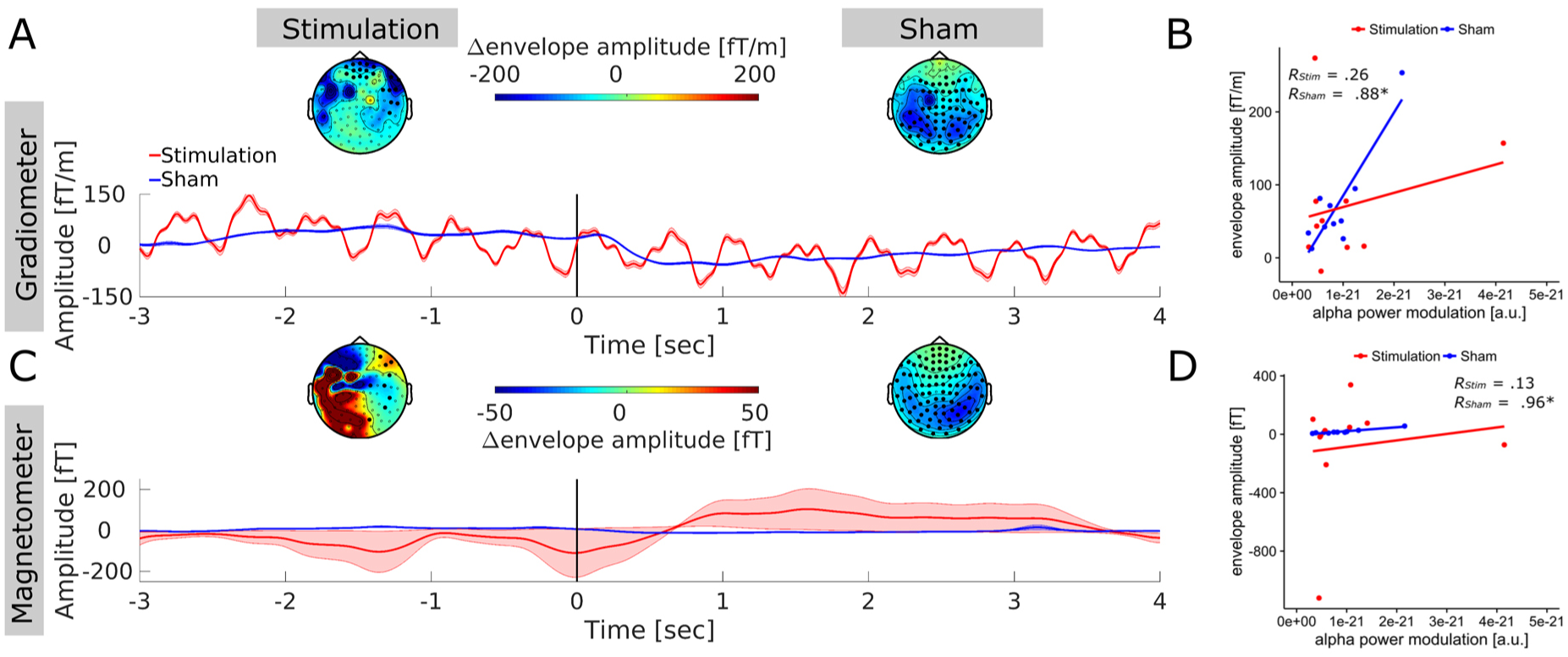
Event-related artifact envelope. **(A)** Topography and time course of the artifact envelope around stimulus onset in gradiometer sensors. Topographies represent the amplitude difference of the envelope around the stimulation frequency between the reference (-2.5 to ‐0.5 sec) and the testing period (0 to 2 sec). Bold sensors mark locations in which this difference was significant. Data of the sham group is depicted for comparison and reflects the task-related modulation of endogenous alpha oscillations (visible shortly after stimulus onset, vertical black bar at 0 sec) as no stimulation artifact was introduced to the data. Envelope epochs of all subjects were demeaned before averaging to enhance comparability of the envelope modulation. Shaded areas depict standard error of the mean (*S.E.M.*). Gradiometer time-courses are strongly dominated by rhythmic modulation around 1 Hz - 2 Hz that potentially reflects a technical artifact in this sensor type. **(B)** Correlation between event-related modulation of the artifact envelope in gradiometer sensors and event-related alpha power modulation within the ROI after beamforming. The absence of a significant (or even moderately high) correlation in the stimulation group provides supporting evidence that the effects observed in source-space are not driven by systematic event-related modulations of tACS artifact strength. **(C)** Topography and time course of the artifact envelope around stimulus onset in magnetometer sensors. **(D)** Correlation between event-related modulation of the artifact envelope in magnetometers and alpha power modulation within ROI after beamforming. Similar to the gradiometer data, no correlation between source-level effects and artifact tACS artifact modulation was observed.

## 4 Discussion

So far, only few studies investigated the effects of tACS on oscillatory activity in the human brain *during* stimulation (Helfrich et al., 2014; Voss et al., 2014; Ruhnau et al., 2016). The current findings add to this line of research by characterizing how event-related oscillatory activity in the brain reacts to externally applied perturbations in the same frequency band during cognitive tasks.

Results show that, rather than counteracting or overwriting the event-related down-regulation of oscillatory power during the MR task, tACS facilitated the pre-existing difference between pre‐ and post-stimulus power in the alpha band. Although this finding converges with observations of facilitated ERD after tACS (Kasten and Herrmann, 2017), it is important to emphasize that online effects of tACS cannot directly be inferred from after-effects. While computational models and animal experiments suggest entrainment as the core mechanism for tACS online effects (Fröhlich and McCormick, 2010; Ozen et al., 2010; Reato et al., 2010), there is increasing evidence that after-effects of tACS might be better explained by mechanisms of neural plasticity (Zaehle et al., 2010; Vossen et al., 2015). Different mechanisms of action could in principle lead to different effects of tACS on event-related oscillations during and after stimulation, rendering direct observations of tACS online effects inevitable to accurately predict and to understand behavioral outcomes in tACS experiments.

The observed enhancement of event-related alpha power modulation explains previous results of better performance in the MR task already during tACS (Kasten and Herrmann, 2017), but contradicts with observations of Neuling et al. (2015). That study reported a tendency for reduced alpha desynchronization elicited by a passive visual task during tACS. However, authors calculated relative change (computed similar to ERD/ERS) to capture event-related alpha desynchronization, which is more vulnerable to residual artifacts in the data. As shown in **Equation 2 and 3**, such a residual leads to a biased (larger) denominator, resulting in systematic underestimations of event-related desynchronization (ERD) within the stimulated frequency band. Using the absolute power difference (here termed ERΔ_Pow_) between two time intervals within the same stimulation condition (i.e. pre-/post-stimulus alpha power) appears to be a more robust measure to capture online effects of tACS. Here, the residual artifact cancels out during the subtraction process. Importantly, this cancelation assumes that the strength of the residual is relatively stable between conditions and uncorrelated with the task. Such systematic modulations could in principle occur if the task elicits systematic changes in physiological processes like heart-beat, respiration, or skin conductance (Noury et al., 2016). While there was no evidence for such a systematic change in artifact strength that could explain the observed pattern in the current data, this fact has to be taken into account e.g. when using stimuli that can elicit stronger physiological responses (i.e. emotional pictures, demanding motor tasks). However, the impact of these modulations on the artifact suppression as compared to the size of the physiological effect on the brain has not been thoroughly characterized yet.

In addition to the observed effect of tACS on power modulations in the alpha band, data revealed a trend towards increased event-related power modulations in the lower beta band during tACS. On the one hand, this observation could be indicative of a rather unspecific effect of tACS (Kleinert et al., 2017), on the other hand the effect in the lower beta band might reflect entrainment or resonance phenomena at the first harmonic of subjects’ stimulation frequency (Herrmann, 2001; Herrmann et al., 2016a).

Contradicting with our previous finding of a prolonged ERD increase in the alpha band after tACS (Kasten and Herrmann, 2017) and despite the massive online effects, only a short-lasting after-effect during the first block after stimulation can be observed, if at all. Several studies successfully showed outlasting effects of tACS on alpha power during rest (i.e. Kasten et al., 2016; Neuling et al., 2013; Veniero et al., 2015; Vossen et al., 2015). A possible explanation for this lack of a sustained outlasting tACS effect might be that stimulation intensity was relatively low as compared to the aforementioned experiments.

Similar to our previous study a significantly stronger increase in mental rotation performance was observed in the stimulation group as compared to sham. Unfortunately, it cannot be ruled out that this effect might have partly been driven by differences in participants’ baseline performance and ceiling effects in the two groups. This could also explain the absence of previously observed correlations between performance increase and facilitated alpha power modulation (Kasten and Herrmann, 2017), that would have further supported the physiological findings. While such ceiling effects would render the current behavioral results uninformative, they do not contradict the physiological effects, which were the main focus of the current study. Mental rotation tasks induce comparably long lasting event related power modulations (Michel et al., 1994), which is a beneficial property when studying tACS effects on event-related oscillations. In the current experiment, this came at the cost of overall high task performance in both groups. Future studies might therefore benefit from more difficult mental rotation paradigms (i.e. only including large rotation angles).

Apart from insights to online effects of tACS on event-related oscillations, the current study made an attempt to quantify the artifact suppression capabilities of LCMV beamforming. To this end, power around the stimulated frequency during tACS was compared to an artifact free estimate of participants’ natural brain signal (alpha power). This allows to estimate the size of the stimulation artifact relative to the brain signal of interest before and after artifact suppression. In the current study, this artifact-to-brain-signal-ratio was suppressed from more than 2,500,000:1 to approximately 3:1, with stronger artifacts around stimulation electrodes and cables (approx. 10:1). Since the power values obtained during stimulation will always contain a mixture of residual tACS artifact and brain signal, this ratio can only provide an upper boundary for the size of the residual artifact. Alpha power increases up to 300-400% fall into a physiologically plausible range for spontaneous alpha power changes or tACS effects, as they are observed i.e. in studies on tACS after-effects (Neuling et al., 2013; Kasten and Herrmann, 2017; Stecher et al., 2017). The artifact-to-brain-signal-ratio might nevertheless be a useful tool for future studies to assess whether a residual artifact falls within the same order of magnitude as the brain signal of interest and could also be used to evaluate and optimize the performance of artifact suppression approaches i.e. by tuning relevant parameters. So far artifact suppression approaches have mostly been evaluated subjectively, i. e. by inspecting raw time series, (time-)frequency spectra or ERPs (Helfrich et al., 2014; Neuling et al., 2015; Witkowski et al., 2016). The artifact-to-brain-signal-ratio provides a more objective evaluation of the artifact size relative to the brain signal of interest, which is scale free and thus also allows easy comparison of different artifact suppression approaches with different measurement modalities (EEG/MEG, LCMV, template subtraction etc.). In addition, the mapping of residual artifact strength allows to check for overlap between ‘hot spots’ of residual artifacts and regions of interest.

The findings presented in the study provide first direct insights to online effects of tACS on event-related oscillations in humans. The effects were investigated using a rather simplistic approach utilizing only two conditions (stimulation vs. sham) and one stimulation frequency targeting posterior alpha oscillations with a Cz-Oz montage. This path was chosen to first establish an analysis and control analysis framework for the investigation of tACS online effects on event-related oscillations, before approaching more complex designs, requiring larger sample sizes and higher computational efforts. TACS experiments generally allow for a multitude of control and contrast conditions including alternative electrode montages and frequencies. The current study can therefore neither resolve frequency nor montage specificity of tACS effects. However, with the present results and the proposed analysis pipeline the current study provides a starting point paving the way to further investigate montage and frequency specificity of tACS effects on event-related oscillatory dynamics during various cognitive tasks.

## 5 Author contributions

FHK, BM and CSH conceived the study and wrote the manuscript. FHK and BM collected the data. FHK analyzed the data. BM and CSH provided equipment and funding for the study.

## Acknowledgements

The authors would like to thank Yvonne Wolf-Rosier for her invaluable efforts and assistance during data collection.

This research was supported by a grant of the German Research Foundation (Deutsche For-schungsgemeinschaft, DFG) awarded to Christoph S. Herrmann (DFG, SPP, 1665 HE 3353/8-2).

## 7 Figure Captions

## Notes

**Conflict of interest:** CSH has filed a patent application on brain stimulation and received honoraria as editor from Elsevier Publishers, Amsterdam. FHK and BM declare no competing interests.

## References

Alekseichuk I, Diers K, Paulus W, Antal A (2016) Transcranial electrical stimulation of the occipital cortex during visual perception modifies the magnitude of BOLD activity: A combined tES‒fMRI approach. Neuroimage 140:110‒117.

Ali MM, Sellers KK, Fröhlich F (2013) Transcranial Alternating Current Stimulation Modulates Large-Scale Cortical Network Activity by Network Resonance. J Neurosci 33:11262‒11275.

Basar E, Basar-Eroglu C, Karakas S, Schürmann M (2000) Brain oscillations in perception and memory. Int J Psychophysiol 35:95‒124.

Brunoni AR, Amadera J, Berbel B, Volz MS, Rizzerio BG, Fregni F (2011) A systematic review on reporting and assessment of adverse effects associated with transcranial direct current stimulation. Int J Neuropsychopharmacol 14:1133‒1145.

Buzsáki G (2006) Rhythms of the Brain. Oxford University Press.

Cabral-Calderin Y, Williams KA, Opitz A, Dechent P, Wilke M (2016) Transcranial alternating current stimulation modulates spontaneous low frequency fluctuations as measured with fMRI. Neuroimage 141:88‒107.

Fröhlich F, McCormick DA (2010) Endogenous Electric Fields May Guide Neocortical Network Activity. Neuron 67:129‒143.

Ganis G, Kievit R (2015) A New Set of Three-Dimensional Shapes for Investigating Mental Rotation Processes: Validation Data and Stimulus Set. J Open Psychol Data 3.

Helfrich RF, Schneider TR, Rach S, Trautmann-Lengsfeld SA, Engel AK, Herrmann CS (2014) Entrainment of Brain Oscillations by Transcranial Alternating Current Stimulation. Curr Biol 24:333‒339.

Herrmann CS (2001) Human EEG responses to 1-100 Hz flicker: resonance phenomena in visual cortex and their potential correlation to cognitive phenomena. Exp brain Res 137:346‒353.

Herrmann CS, Murray MM, Ionta S, Hutt A, Lefebvre J (2016a) Shaping Intrinsic Neural Oscillations with Periodic Stimulation. J Neurosci 36:5328‒5337.

Herrmann CS, Strüber D, Helfrich RF, Engel AK (2016b) EEG oscillations: From correlation to causality. Int J Psychophysiol 103:12‒21.

Kar K, Duijnhouwer J, Krekelberg B (2017) Transcranial Alternating Current Stimulation Attenuates Neuronal Adaptation. J Neurosci 37:2325‒2335.

Kar K, Krekelberg B (2014) Transcranial Alternating Current Stimulation Attenuates Visual Motion Adaptation. J Neurosci 34:7334‒7340.

Kasten FH, Dowsett J, Herrmann CS (2016) Sustained Aftereffect of α-tACS Lasts Up to 70 min after Stimulation. Front Hum Neurosci 10:1‒9.

Kasten FH, Herrmann CS (2017) Transcranial Alternating Current Stimulation (tACS) Enhances Mental Rotation Performance during and after Stimulation. Front Hum Neurosci 11:1‒16.

Kleiner M, Brainard DH, Pelli DG, Broussard C, Wolf T, Niehorster D (2007) What’s new in Psychtoolbox-3? Perception 36:S14.

Kleinert M-L, Szymanski C, Müller V (2017) Frequency-Unspecific Effects of θ-tACS Related to a Visuospatial Working Memory Task. Front Hum Neurosci 11:1‒16.

Klimesch W (1999) EEG alpha and theta oscillations reflect cognitive and memory performance: A review and analysis. Brain Res Rev 29:169‒195.

Klimesch W, Sauseng P, Gerloff C (2003) Enhancing cognitive performance with repetitive transcranial magnetic stimulation at human individual alpha frequency. Eur J Neurosci 17:1129‒1133.

Klimesch W, Sauseng P, Hanslmayr S (2007) EEG alpha oscillations: The inhibition-timing hypothesis. Brain Res Rev 53:63‒88.

Lustenberger C, Boyle MR, Foulser AA, Mellin JM, Fröhlich F (2015) Functional role of frontal alpha oscillations in creativity. Cortex 67:74‒82.

Mäkelä N, Sarvas J, Ilmoniemi RJ (2017) Proceedings #17. A simple reason why beamformer may (not) remove the tACS-induced artifact in MEG. Brain Stimul 10:e66-e67.

Marshall L, Helgadóttir H, Mölle M, Born J (2006) Boosting slow oscillations during sleep potentiates memory. Nature 444:610‒613.

Michel CM, Kaufman L, Williamson SJ (1994) Duration of EEG and MEG α Suppression Increases with Angle in a Mental Rotation Task. J Cogn Neurosci 6:139‒150.

Neuling T, Rach S, Herrmann CS (2013) Orchestrating neuronal networks: sustained aftereffects of transcranial alternating current stimulation depend upon brain states. Front Hum Neurosci 7:161.

Neuling T, Ruhnau P, Fuscà M, Demarchi G, Herrmann CS, Weisz N (2015) Friends, not foes: Magnetoencephalography as a tool to uncover brain dynamics during transcranial alternating current stimulation. Neuroimage 118:406‒413.

Neuling T, Ruhnau P, Weisz N, Herrmann CS, Demarchi G (2017) Faith and oscillations recovered: On analyzing EEG/MEG signals during tACS. Neuroimage 147:960‒963.

Nolte G (2003) The magnetic lead field theorem in the quasi-static approximation and its use for magnetoencephalography forward calculation in realistic volume conductors. Phys Med Biol 48:3637‒3652.

Noury N, Hipp JF, Siegel M (2016) Physiological processes non-linearly affect electrophysiological recordings during transcranial electric stimulation. Neuroimage 140:99‒109.

Noury N, Siegel M (2017) Phase properties of transcranial electrical stimulation artifacts in electrophysiological recordings. Neuroimage 158:406‒416.

Oldfield RC (1971) The assessment and analysis of handedness: The Edinburgh inventory. Neuropsychologia 9:97‒113.

Oostenveld R, Fries P, Maris E, Schoffelen JM (2011) FieldTrip: Open source software for advanced analysis of MEG, EEG, and invasive electrophysiological data. Comput Intell Neurosci 2011:1‒9.

Ozen S, Sirota A, Belluscio MA, Anastassiou CA, Stark E, Koch C, Buzsaki G (2010) Transcranial Electric Stimulation Entrains Cortical Neuronal Populations in Rats. J Neurosci 30:11476‒11485.

Pfurtscheller G, Lopes Da Silva FH (1999) Event-related EEG/MEG synchronization and desynchronization: Basic principles. Clin Neurophysiol 110:1842‒1857.

Reato D, Rahman A, Bikson M, Parra LC (2010) Low-Intensity Electrical Stimulation Affects Network Dynamics by Modulating Population Rate and Spike Timing. J Neurosci 30:15067‒15079.

Ruhnau P, Neuling T, Fuscá M, Herrmann CS, Demarchi G, Weisz N (2016) Eyes wide shut: Transcranial alternating current stimulation drives alpha rhythm in a state dependent manner. Sci Rep 6:27138.

Shepard RN, Metzler J (1971) Mental rotation of three-dimensional objects. Science 171:701‒703.

Soekadar SR, Witkowski M, Cossio EG, Birbaumer N, Robinson SE, Cohen LG (2013) In vivo assessment of human brain oscillations during application of transcranial electric currents. Nat Commun 4:2032.

Stecher HI, Pollok TM, Strüber D, Sobotka F, Herrmann CS (2017) Ten Minutes of α-tACS and Ambient Illumination Independently Modulate EEG α-Power. Front Hum Neurosci 11:1‒10.

Taulu S, Simola J, Kajola M (2005) Applications of the signal space separation method. IEEE Trans Signal Process 53:3359‒3372.

Van Veen BD, Van Drongelen W, Yuchtman M, Suzuki A (1997) Localization of brain electrical activity via linearly constrained minimum variance spatial filtering. IEEE Trans Biomed Eng 44:867‒880.

Veniero D, Vossen A, Gross J, Thut G (2015) Lasting EEG/MEG Aftereffects of Rhythmic Transcranial Brain Stimulation: Level of Control Over Oscillatory Network Activity. Front Cell Neurosci 9:477.

Violante IR, Li LM, Carmichael DW, Lorenz R, Leech R, Hampshire A, Rothwell JC, Sharp DJ (2017) Externally induced frontoparietal synchronization modulates network dynamics and enhances working memory performance. Elife 6.

Voss U, Holzmann R, Hobson A, Paulus W, Koppehele-Gossel J, Klimke A, Nitsche M a (2014) Induction of self awareness in dreams through frontal low current stimulation of gamma activity. Nat Neurosci 17:810‒812.

Vossen A, Gross J, Thut G (2015) Alpha Power Increase After Transcranial Alternating Current Stimulation at Alpha Frequency (α-tACS) Reflects Plastic Changes Rather Than Entrainment. Brain Stimul 8:499‒508.

Vosskuhl J, Huster RJ, Herrmann CS (2016) BOLD signal effects of transcranial alternating current stimulation (tACS) in the alpha range: A concurrent tACS‒fMRI study. Neuroimage 140:118‒125.

Wach C, Krause V, Moliadze V, Paulus W, Schnitzler A, Pollok B (2013) Effects of 10Hz and 20Hz transcranial alternating current stimulation (tACS) on motor functions and motor cortical excitability. Behav Brain Res 241:1‒6.

Witkowski M, Garcia-Cossio E, Chander BS, Braun C, Birbaumer N, Robinson SE, Soekadar SR (2016) Mapping entrained brain oscillations during transcranial alternating current stimulation (tACS). Neuroimage 140:89‒98.

Zaehle T, Rach S, Herrmann CS (2010) Transcranial Alternating Current Stimulation Enhances Individual Alpha Activity in Human EEG. PLoS One 5:13766.

